# Transcriptome Enhanced Rice Grain Metabolic Model Identifies Histidine Level as a Marker for Grain Chalkiness

**DOI:** 10.1101/2024.10.14.618272

**Authors:** Niaz Bahar Chowdhury, Anil Kumar Nalini Chandran, Harkamal Walia, Rajib Saha

## Abstract

Rising temperatures due to global warming can negatively impact rice grain quality and yield. This study investigates the effects of increased warmer night temperatures (WNT), a consequence of global warming, on the quality of rice kernel, particularly grain chalkiness. By integrating computational and experimental approaches, we used a rice grain metabolic network to discover the metabolic factors of chalkiness. For this, we reconstructed the rice grain genome-scale metabolic model (GSM), iOSA3474-G and incorporated transcriptomics data from three different times of the day (dawn, dawn 7h, and dusk) for both control and WNT conditions with iOSA3474-G. Three distinct growth phases: anoxia, normoxia, and hyperoxia, were identified in rice kernels from the GSMs, highlighting the grain-filling pattern under varying oxygen levels. We predicted histidine as a marker of normoxia, during which kernel chalkiness occurs. Moreover, we proposed tyrosine as a marker for the hyperoxic growth phase. We also proposed a potential link between monodehydroascorbate reductase, an enzyme with evolutionary significance dating back to the carboniferous era, in regulating the hyperoxic growth phase. Metabolic bottleneck analysis identified nucleoside diphosphate kinase as a central regulator of metabolic flux under different conditions. These findings provide targeted insights into the complex metabolic network governing rice grain chalkiness under WNT conditions. Integration of GSM and transcriptomics data, enhanced our understanding of the intricate relationship between environmental factors, metabolic processes, and grain quality and also offer markers that can be useful to develop rice with improved resilience.

## INTRODUCTION

Rice is a staple for more than half of the world’s population, supplying over 21% of total caloric demands (Yuan *et al*., 2021). With projected increase in global population to 9 billion by 2050 (Roberts, 2011), the demand for rice will continue to rise sharply. However, global warming poses a significant challenge to meeting this demand. Predictions suggest a rise in global mean surface air temperatures of 1.0–3.7°C by 2100 (He *et al*., 2018), placing rice production cycle at heightened risk, particularly during grain filling, due to heat stress (Lyman *et al*., 2013). Recent studies also underscore the asymmetric increase in night-time temperatures compared to day-time temperatures (Sillmann *et al*., 2014), exerting an additional vulnerability to rice grain yield.

Exposure to warmer night-time (WNT) conditions during grain filling significantly reduces pollen viability, heightens spikelet sterility, compromises membrane integrity, and hinders grain growth, ultimately resulting in poor seed-set and reduced individual grain weight (Mohammed and Tarpley, 2010; Coast *et al*., 2016; Cao *et al*., 2017). Changes in individual grain weight are often linked to limited carbohydrate supply (Nakamura *et al*., 1989) and alterations in starch metabolism enzymes (Bahuguna *et al*., 2017). Besides reduced grain yield, exposure to WNT also adversely affects grain quality, which often manifests as increased chalkiness (Lanning *et al*., 2011). Chalkiness is the opaque part of the grain found in an otherwise translucent white endosperm of rice. Chalk formation in rice grain arises from loosely packed starch granules, resulting in air spaces between amyloplasts (Ashida *et al*., 2009), resulting in a higher proportion of broken grains and a significant decrease in the economic value of the rice (Zhao and Fitzgerald, 2013).

Despite some understanding of rice grain chalkiness, further insights into its implications on the overall rice grain metabolism remains scarce. To address this, the development of a genome-scale metabolic model (GSM) (Alsiyabi *et al*., 2022) specific to rice grain, coupled with transcriptome data that potentially captures the increased chalkiness under WNT, could provide a novel insights into the role of metabolic perturbance under WNT. Previous successes in integrating transcriptomics data with GSMs have been demonstrated in predicting phenotypes, such as maize root phenotype under nitrogen starvation conditions and whole-plant phenotype of maize under heat and cold stress conditions (Chowdhury, Schroeder, *et al*., 2022; Chowdhury *et al*., 2023). Therefore, this approach holds promise for elucidating the metabolic pathways associated with rice grain chalkiness. While previous rice GSMs exist, they are consolidated whole-plant models and primarily focus on the photosynthetic apparatus under various conditions (Lakshmanan *et al*., 2013; Poolman *et al*., 2013; Chatterjee *et al*., 2017). Although a multi-organ rice metabolic model has been developed recently (Shaw and Cheung, 2021), it lacks the resolution required to capture grain-related metabolism. Furthermore, these studies have not integrated relevant transcriptomics data with the models, a crucial process known as contextualization, essential for accurately predicting phenotypes (Chowdhury, Schroeder, *et al*., 2022). Thus, the development of a grain-specific rice model, contextualized with a transcriptome response that is known to increase grain chalkiness under WNT stress, is essential for comprehensively understand the system-wide metabolic impacts of grain chalkiness.

To explore rice grain chalkiness, we constructed the first-ever rice grain GSM, iOSA3474-G, and contextualized it with a previously published WNT transcriptomics dataset (Desai *et al*., 2021) across three time points (dawn, dawn 7h, and dusk) for both control and WNT conditions using the EXTREAM algorithm (Chowdhury *et al*., 2023) that we have recently developed and accurately predicted the phenotype of maize plant under heat and cold stress conditions. Our analysis identified three distinct growth phases - anoxia, normoxia, and hyperoxia - in rice grains from the GSMs, elucidating the grain-filling dynamics in the context of varying oxygen levels.

Notably, we predicted histidine as a metabolic marker for normoxia, the phase coinciding with grain chalkiness occurrence, while tyrosine as a metabolic marker for the hyperoxic growth phase. Furthermore, our investigation predicted the significant role of monodehydroascorbate reductase (MDAR), an enzyme with evolutionary significance tracing back to the carboniferous era, in regulating the hyperoxic growth phase. Through metabolic bottleneck analysis, we proposed nucleoside diphosphate kinase as a central regulator of metabolic flux under both different conditions. These findings offer targeted insights into the intricate metabolic network governing rice grain chalkiness under WNT conditions. By integrating GSM and transcriptomics data, this approach not only advances our comprehension of the complex interplay between environmental factors, metabolic pathways, and crop quality but also presents practical avenues for enhancing crop resilience to global warming.

## RESULTS AND DISCUSSION

### Reconstruction of Genome-Scale Metabolic Model of Rice Grain

Maize, being a monocot, shares significant common ancestry with rice (Murat *et al*., 2017). Comparative studies of Quantitative Trait Loci (QTL) between these two species have revealed that comparable traits are typically governed by QTLs located within syntenic regions (Paterson *et al*., 1995). Furthermore, genes influencing grain shape and weight in rice, such as GS3, GW2, and GS5, have been demonstrated to regulate similar traits in maize (Li *et al*., 2011). Therefore, we choose to use our recently published maize kernel model (Chowdhury *et al*., 2023) as a blueprint to reconstruct the rice grain model. Figure 1a shows the rice grain metabolic model reconstruction workflow.

**Fig. 1:**
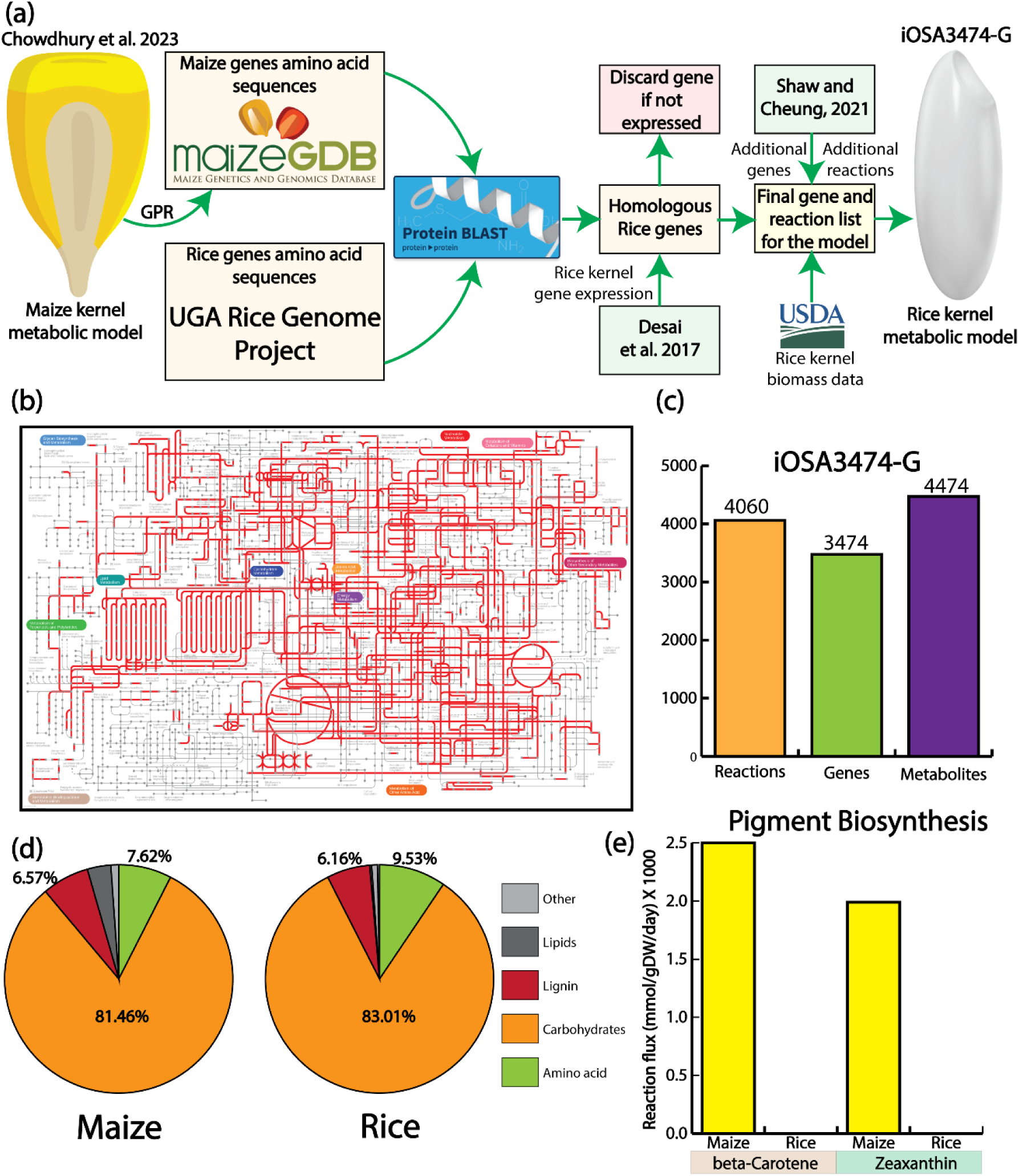
Genome-scale metabolic model reconstruction of rice grain. (a) iOSA343474-G reconstruction through homology search against maize kernel metabolic model, literature search, and existing whole plant rice metabolic model. Biomass equation was reconstructed from the data obtained from USDA. (b) A metabolic network of iOSA3474-G generated from KEGG. (c) Statistic of the iOSA3474-G. (d) Biomass constituent comparison between maize kernel and rice grain. (e) iOSA3474-G cannot produce major pigments, beta-carotene and zeaxanthin, which maize kernel can produce.

Fig. 1b shows the metabolic map of rice grain model. Overall, the model, iOSA3474-G, has 3474 genes, 4060 reactions, and 4474 metabolites (Fig. 1 c) and is the most comprehensive rice grain metabolic model till date.

Next, rice grain biomass composition data was collected from the USDA website (Supplementary Data S1) and formulated in a way so that the molecular weight of biomass is 1 mmol/gDW (Chan *et al*., 2017). Biomass composition comparison between maize kernel and rice grain revealed the difference in protein, lipid, fatty acids, and pigments compositions. The mole percentage of carbohydrate in rice is higher than the maize kernel (Fig. 1 d). Interestingly, amino acid composition in rice grain is also higher than the maize kernel. Contrastingly, lipid content is maize is higher (3.41%) in maize kernel compared to the rice grain (0.24%).

One of the key differences between maize kernel and rice grain is its inability to produce pigments, without genetic perturbations (Zhao *et al*., 2022). Contrary to that, p-carotene and zeaxanthin are the major pigments in the maize kernel (Song *et al*., 2016). Interestingly, iOSA3474-G predicted no flux through p-carotene and zeaxanthin biosynthesis pathway (Fig. 1 e). However, the maize kernel model was able to biosynthesize both the pigments (Fig. 1 e). Thereby, iOSA3474-G was able to capture biologically meaningful phenotype of rice grain.

### Connecting Warmer Night Temperature (WNT) with Grain Chalkiness

As iOSA3474-G was able to predict biologically relevant phenotypes for the rice grain, we used this model to gain further insights on rice grain metabolism under WNT. For that, we used a published transcriptomic dataset for Dawn control/WNT, Dawn 7h control/WNT, and Dusk control/WNT (Desai *et al*., 2021). These transcriptomic data showed high degree of Pearson correlation (Fig. 2 a). The lowest degree of correlation was found between Dawn control-Dawn 7h control, Dawn WNT-Dawn 7h WNT conditions, which was 0.79. This indicated 7h after dawn, a transcriptomic response changes compared to the dawn conditions. This finding is consistent with the previous study (Desai *et al*., 2021), which reported higher activity of photosynthesis machinery at Dawn 7h compared to the Dawn condition.

**Fig. 2:**
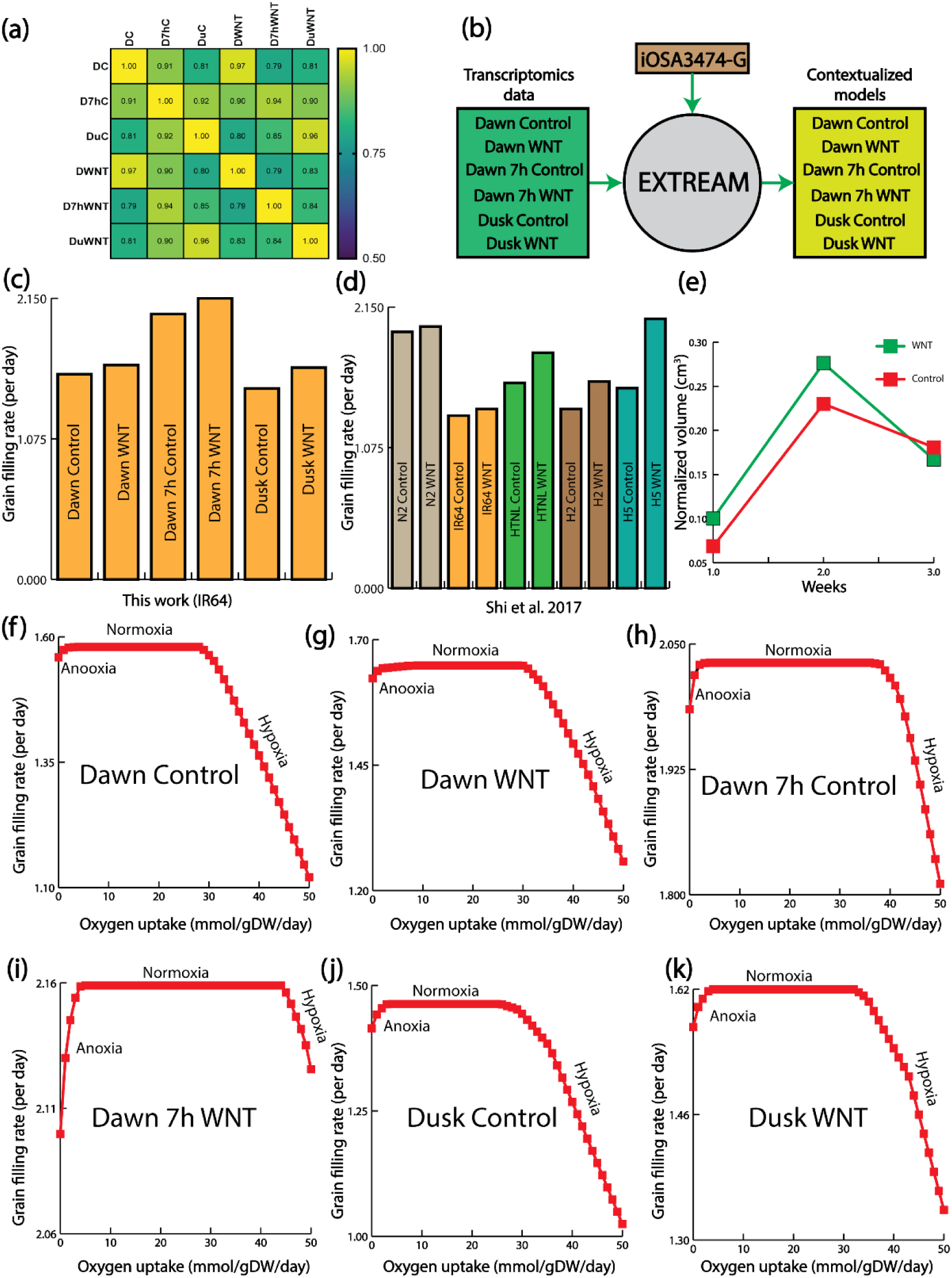
Warmer night temperature promotes gran chalkiness. (a) Dawn control/Dawn WNT, Dawn 7h control/Dawn 7h WNT, and Dusk control/Dusk WNT are strongly correlated, indicating a similar transcriptomic response between control and WNT conditions. (b) EXTREAM was used to reconstruct six contextualized models using transcriptomics data and iOSA3474-G. (c) Contextualized iOSA3474 predicted grain filling rates. (d) Grain filling rates from literature for different rice genotypes. (e) Experimentally measured grain volume data confirmed higher grain filling rate in WNT than control condition. (f) Grain filling rate with varying oxygen uptake for Dawn control. (g) Grain filling rate with varying oxygen uptake for Dawn WNT. (h) Grain filling rate with varying oxygen uptake for Dawn 7h control. (i) Grain filling rate with varying oxygen uptake for Dawn 7h WNT. (j) Grain filling rate with varying oxygen uptake for Dusk control. (k) Grain filling rate with varying oxygen uptake for Dusk WNT.

To project these transcriptomics data into the fluxomics space, we incorporated these transcriptomics data with iOSA3474-G using our recently developed tool called EXTREAM (Chowdhury *et al*., 2023) (Fig. 2 b). Rice panicle transcriptomics data is particularly well-suited to understand rice grain metabolism, as the panicle itself contains the developing grains and acts as the central conduit for nutrient transfer and metabolic regulation throughout grain maturation. Key metabolic activities within the panicle—such as starch synthesis, lipid metabolism, and amino acid production—are crucial for grain development, providing essential precursors and controlling the energy dynamics necessary for grain filling. Because the grains are an integral part of the panicle, its gene expression profiles inherently reflect the metabolic processes occurring within the grains, thereby offering deeper insights into the pathways that determine grain quality and yield. EXTREAM previously predicted correct phenotypes for whole maize plant under control and temperature stress conditions and uncovered the role of energy production and reducing power generation in explaining metabolic bottlenecks of temperature stress conditions (Chowdhury *et al*., 2023). EXTREAM returned six contextualized models which were further probed to understand metabolic underpinnings of WNT conditions. Contextualized models predicted higher grain filling rates for WNT conditions (Fig. 2 c). Previous studies (Shi *et al*., 2017) also predicted this pattern, not only for IR64, but also for other rice genotypes (Fig. 2 d). *De novo* experimental data, calculating grain volume for control and WNT conditions, also supported the higher grain filling rate in the WNT conditions (Fig. 2 e). Grain weight is determined by a balance between grain-filling rate and duration of grain filling. Although WNT conditions consistently showed higher grain filling rates, it was reported that, due to lack of assimilates (Kobata and Uemuki, 2004) and loss of sink activity (Kim *et al*., 2011), an early termination of grain filling in rice can occur under higher temperature. Moreover, photosynthesis, transpiration, and respiration are temperature-sensitive processes that contribute to grain weight. However, only respiration occurs consistently during the day and night. An increase in nighttime respiration has been associated with high nighttime temperatures. Therefore, increase in dark respiration is often considered as the primary mechanism for the observed high grain filling rate and decreased grain-filling duration (Desai *et al*., 2021).

In WNT, the density of oxygen is reduced in the atmosphere, thus less oxygen is available for the grain. Therefore, we asked if the grain growth is changing with varying oxygen uptake. We plotted growth profile of grain in all six different conditions, and we recognized three different growth regions: anoxia (oxygen deficient condition), normoxia (normal oxygen level), and hyperoxia (excess oxygen level). A similar growth pattern was observed for the barley seed (Grafahrend-Belau *et al*., 2009). However, for the barley seed, the normoxia phase was brief. Thus, this study, for the first time, reported three distinct growth phases of grain development. Previous works suggested that WNT induces chalkiness in rice grain (Shi *et al*., 2017) (Supplementary Fig. S1) and we speculated that because of less oxygen availability during WNT conditions, chalkiness happens early in the normoxia. Thus, characterizing the normoxia growth will reveal metabolic underpinnings of grain chalkiness.

### Histidine as the Potential Metabolic Marker of Normoxia and Grain Chalkiness

As chalkiness is more relevant in the early stage of normoxia, further characterization of early normoxia is required to understand relationship between oxygen availability and chalkiness. As all the three growth phases are readily visible in the Dawn 7h WNT conditions (Fig. 2 i), initial analysis for normoxia was conducted based on that condition. Some plants accelerate glycolysis under anoxia, a mechanism known as the ‘Pasteur effect’. This helps to alleviate the large reduction of energy produced by fermentation as compared with oxidative phosphorylation, especially during the early period of acclimation to anoxia (Greenway and Gibbs, 2003). Thus, under anoxia, grain should performs more substrate level phosphorylation compared to when more oxygen is available. iOSA3474-G interestingly predicted higher substrate level phosphorylation under early normoxia (Fig. 3 a). As energy generation is restricted in the early normoxia, it is likely that subsequent catabolism of different pathways will also be restricted. That will be more applicable to lipid biosynthesis as rice grain has 0.24% lipid, contrary to the 3.4% lipid in maize kernel (Fig. 1 d). Interestingly, similar to the substrate level phosphorylation, iOSA3474-G correctly predicted lower glycerolipid biosynthesis in early normoxia (Fig. 3 b).

**Fig. 3:**
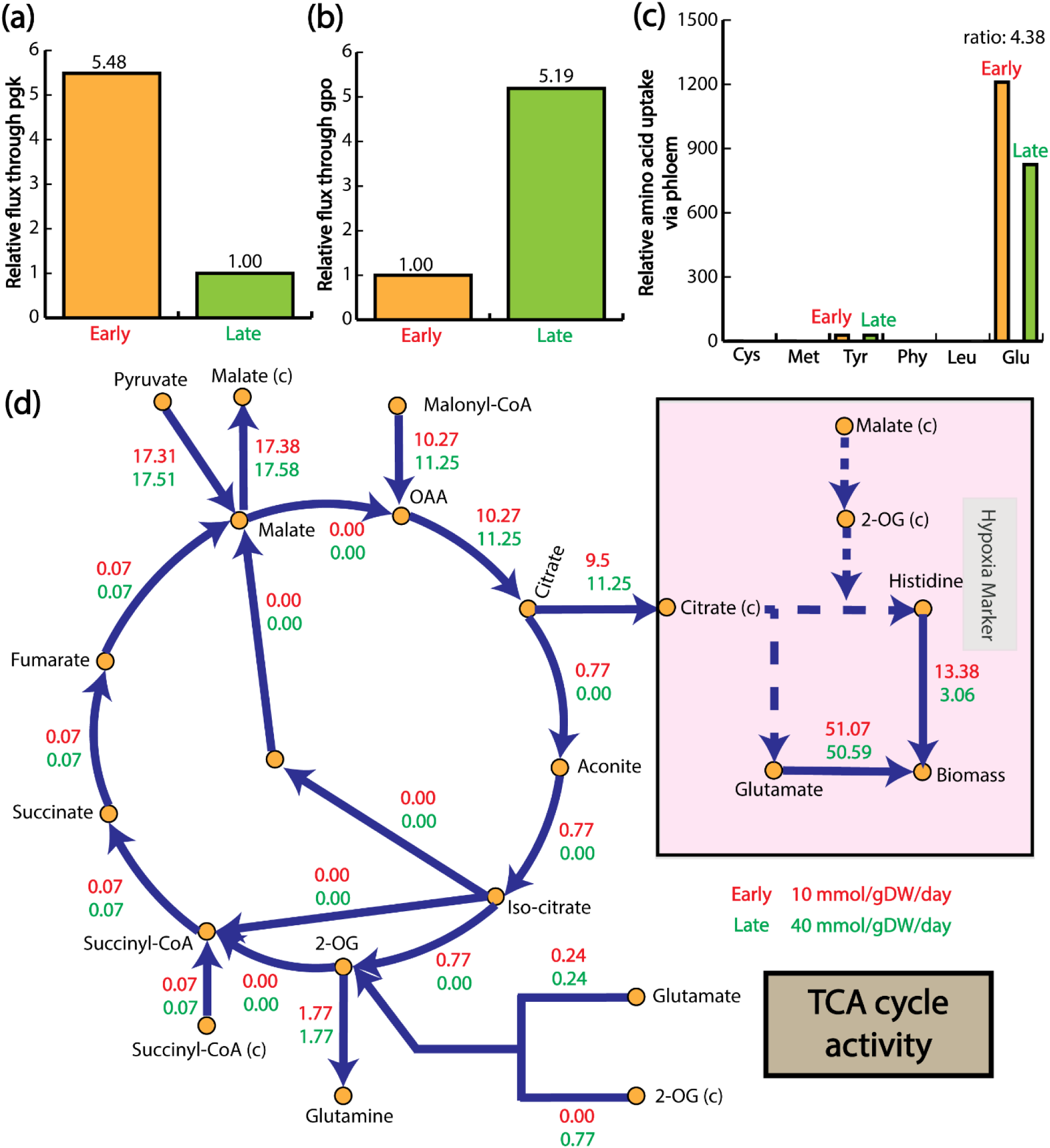
Metabolic characteristic of normoxia. (a) Reduced substrate level phosphorylation. (b) Increased glycerolipid metabolism. (c) High phloem uptake of glutamine by the grain. (d) Metabolic network elucidating the role of histidine as a metabolic marker in grain chalkiness.

Grains serve as a sink tissue, which receive amino acids from the other part of the plant such as root, shoot, and leaf (Egli and Bruening, 2001). Therefore, we aimed to characterize the amino acids that are going to the grain through phloem. Compared to the other amino acids, during normoxia, we noticed very high glutamine uptake by the grain (Fig. 3 c). As one of the major forms of nitrogen in rice (Fukumorita and Chino, 1982), glutamine may have other functions in producing other amino acids (glutamate), TCA components (α-ketoglutarate), and nucleotides (AMP, purines, and pyrimidines), along with the activation of the chaperone function (mediated by HSP response) and antioxidant defence (Cruzat *et al*., 2018). However, how high glutamate uptake through phloem is converted to biomass through other amino acids is still an unanswered question. Therefore, we used iOSA3474-G to check what is the conversion route of the accumulating glutamine to other amino acid that contributed to the biomass growth.

We tracked the high uptake of glutamine in the metabolic network and found histidine to be a metabolic marker for the early normoxia, which represents chalkiness (13.38 mmol/gDW/day of flux towards biomass growth rate in the early normoxia, where as 3.06 mmol/gDW/day of flux towards biomass growth rate in the late normoxia) (Fig. 3 d). Furthermore, tracking glutamate uptake revealed interesting pattern in the TCA cycle. In both early and late normoxia, glyoxylate shunt was turned off. Furthermore, TCA cycle was partially active and produced different metabolites through transport from other compartments (2-OG, glutamate, succinyl-CoA) or by catabolizing higher carbon sources (malonyl-CoA). Besides, it exports metabolites to cytosol as well (malate and citrate). Eventually, the citrate converts to histidine that aids in biomass production. Compared to the late normoxia, early normoxia contributes 4.38 times more histidine to the biomass (Fig. 2 d). Thus, histidine can be a metabolic marker to indicate chalkiness in the rice grain. Previous studies suggested higher histidine level in the less oxygenated condition, supporting the finding of this work (Zhang *et al*., 2021). For other WNT conditions, histidine also showed a similar contribution to the biomass and served as the metabolic marker.

### Tyrosine as the Potential Metabolic Marker of Hyperoxia and the Evolutionary Role of MDAR

We next analyzed the hyperoxic phase, characterized by availability of excess oxygen for the grain. Hyperoxic growth region is not a theoretical prediction, as it has been shown in the lab setting for maize kernel that higher partial pressure of oxygen reduced the overall kernel size (Langer *et al*., 2023). From a previous study (Langer *et al*., 2023), hyperoxia had a modest impact on the number of responsive maize kernel genes, but caused a more rapid rate of kernel development. The most prominently downregulated gene under hyperoxia was a BAX inhibitor-1 (Watanabe and Lam, 2009). In Arabidopsis, this gene suppresses programed cell death. The downregulation of this inhibitor in maize kernels could thus lead to an earlier onset of programed cell death, a characteristic feature of maturing endosperm. Another marker of more rapid kernel development under hyperoxia was the upregulation of an α-zein 3 storage protein typically expressed during late stages of maturation (Stelpflug *et al*., 2016).

In the late hyperoxic growth phase, unlike glutamine in normoxia, grain uptakes 6.67 times higher amount of phenylalanine through phloem tissues compared to the early phase of hyperoxic growth phase (Fig. 4 a). We tracked this high uptake of phenylalanine in the metabolic network and found much higher contribution of tyrosine to the biomass in the late hyperoxia compared to the early hyperoxia (Fig. 4 b). Previous study also supported elevated amount of tyrosine under comparatively high oxygen level (Zhang *et al*., 2021).

**Fig. 4:**
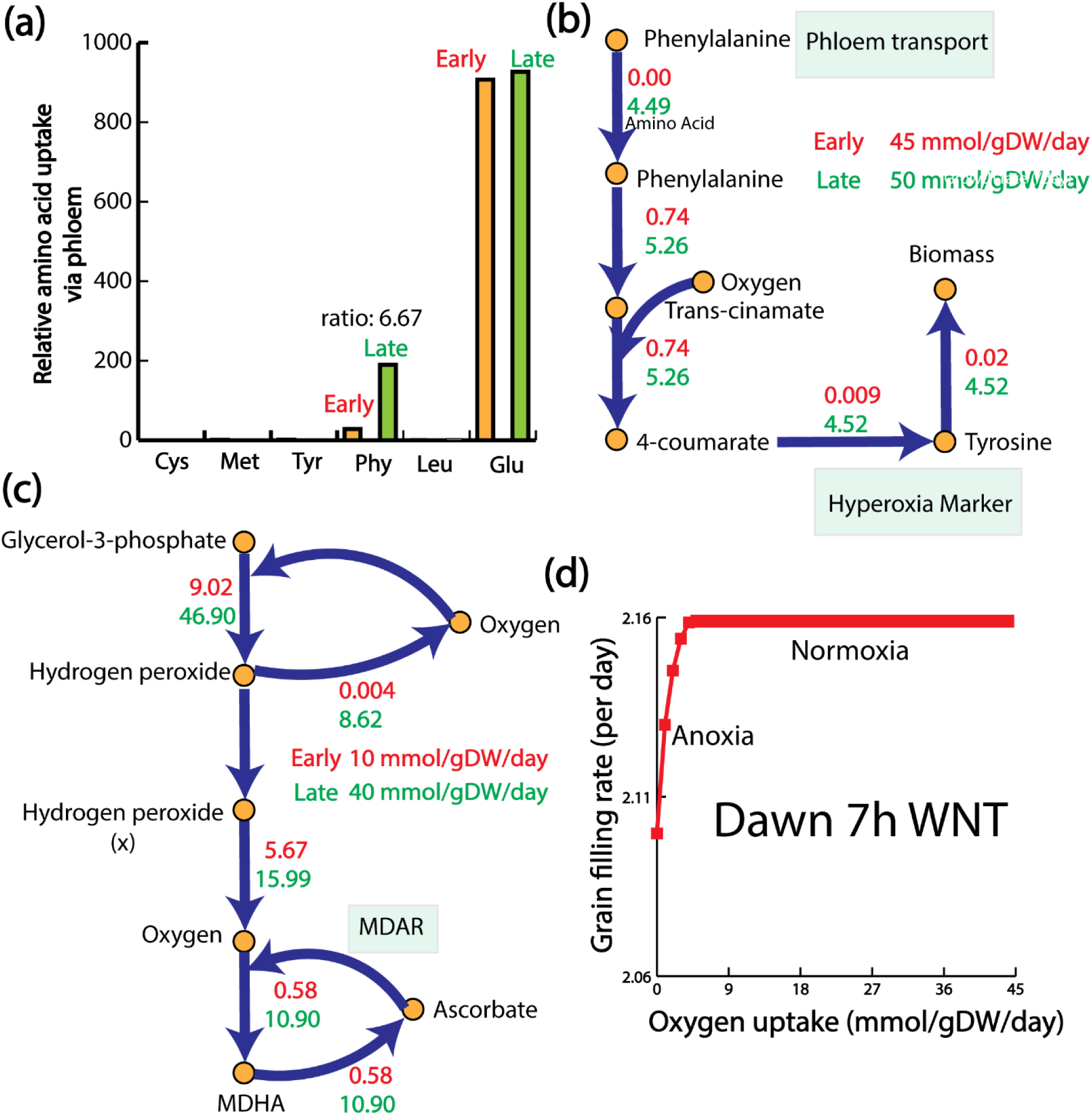
Metabolic characteristic of hyperoxia and the evolutionary role of MDAR. (a) High phloem uptake of phenylalanine by the grain. (b) Metabolic network elucidating the role of tyrosine as a metabolic marker in hyperoxia. (c) Excess oxygen is being processed by monodehydroascorbate reductase (MDAR). (d) If MDAR is knocked out, hyperoxic phase no longer exist. (e) Evolutionary divergence of angiosperms and gymnosperms from spermatophytes.

We also wanted to check where the excess oxygen is being used in the metabolic network in the hyperoxic phase of growth, to understand the grain’s ability to use oxygen. We observed glycerol-3-phosphate being used to produce hydrogen per oxide, which recycled back to oxygen to produce monodehydroascorbate (MDAR). The MDAR recycled back to oxygen with an ascorbate intermediate (Fig. 4 c). This step is catalyzed by an enzyme, monodehydroascorbate reductase (MDAR). MDAR plays a key role in stress tolerance and ascorbate metabolism in plants. MDAR activities have been reported in algae (Haghjou *et al*., 2009), bryophytes (Lunde *et al*., 2006) and higher order plants (Leterrier *et al*., 2005). Moreover, MDAR is one of the key anti-oxidant enzymes responsible for scavenging reactive oxygen species (Sudan *et al*., 2015) and supports the use of excess oxygen by grain through MDAR. When we turn off the MDAR reaction in iOSA3474-G, the hyperoxic phase no longer existed (Fig. 4 d), indicating an essential role of MDAR in scavenging excess oxygen. From an evolutionary perspective, during the carboniferous era (300 to 35 million years ago), atmospheric oxygen levels was 35% (Beerling and Berner, 2000). Spermatophytes encompass the angiosperms and the gymnosperms, whose seeds are not enclosed in an ovary. The two groups diverged around 300 million years ago during the carboniferous era (Savard *et al*., 1994). Thus, to survive the excess oxygen of carboniferous era, plant evolved the metabolic pathways involving MDAR. However, currently, with much lower current oxygen content in the atmosphere (21%), it is plausible that the MDAR activity in the rice grain is a remnant trait from the carboniferous era.

### Potential Role of Nucleotide Diphosphate Kinase and Enoyl-CoA Hydratase in WNT

With a comprehensive understanding of the various growth stages of rice grains in the WNT, next we implemented Metabolic Bottleneck Analysis (MBA) (Chowdhury *et al*., 2023) on six different contextualized models to see how the metabolic bottlenecks changed dynamically in WNT conditions and impacted grain chalkiness. The MBA algorithm expands the flux space of each reaction in a GSM and assess its impact on the biomass growth rate, thereby returns bottleneck reactions. However, we now extended the MBA to return tentative bottleneck genes as well, using the gene-protein-reaction (GPR) association of bottleneck reactions (Fig. 5 a). In this work, we identified 416, 409, 411, 432, 290, and 443 tentative bottleneck genes in Dawn control, Dawn WNT, Dawn 7h control, Dawn 7h WNT, Dusk controļ and Dusk WNT, respectively (Supplementary Fig. S2). Details of all the genes can be found in the Supplementary Data S2. We also performed gene ontology analysis of control and WNT conditions. For all the control conditions, bottleneck genes were pertinent to cellular response to oxidative stress, response to oxygen-containing compounds, phosphorylation, organonitrogen compound metabolic process, etc. (Supplementary Fig. S3). This indicated that, all the control conditions have similar kind of bottlenecks and behaved similarly. Correlation of transcriptomics data (Fig. 2 a) for all the control conditions also supports this (lowest correlation 0.81, between Dawn control and Dusk control conditions).

**Fig. 5:**
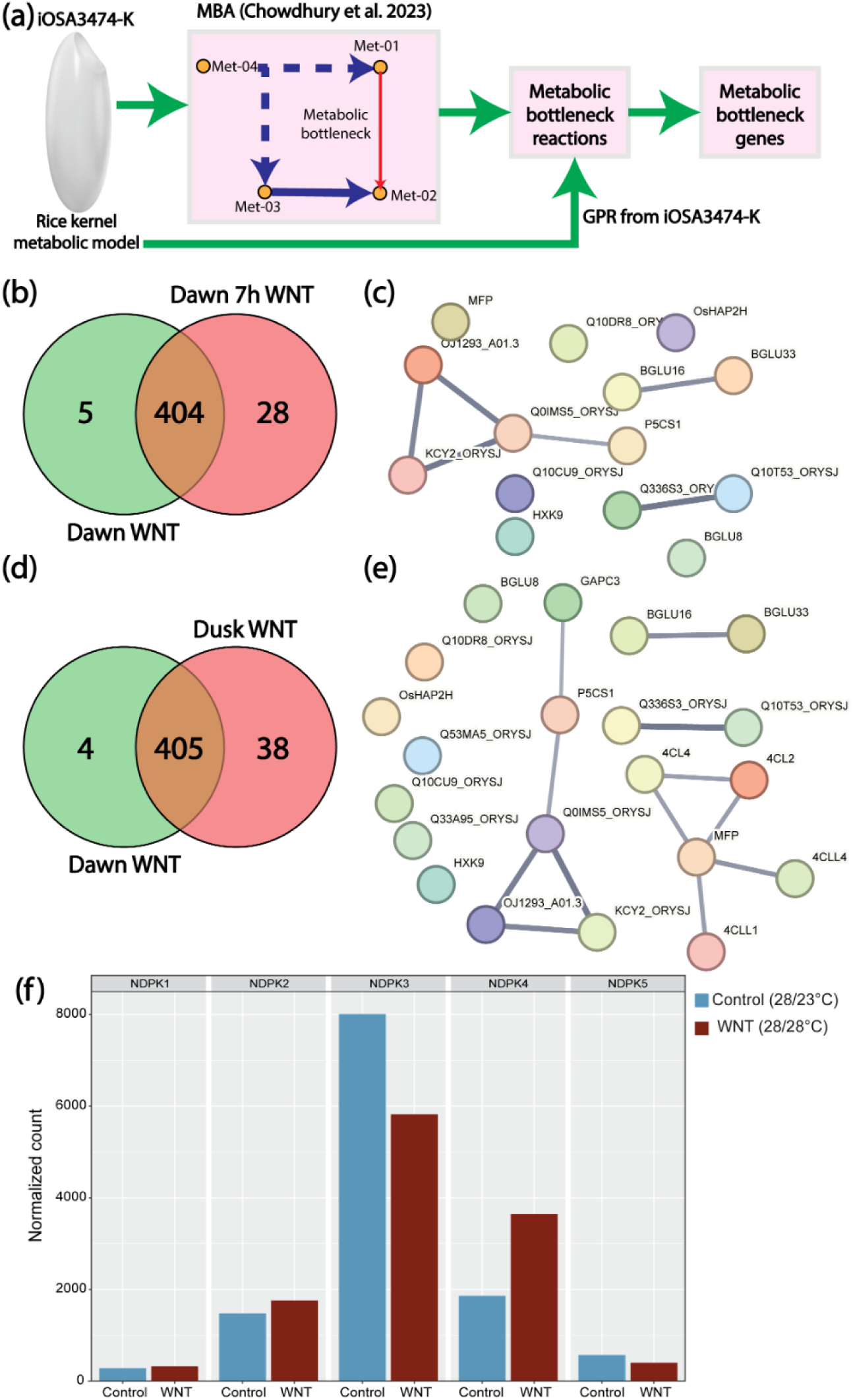
Metabolic Bottlenecks Analysis (MBA) revealed the role of Nucleotide Diphosphate Kinase and Enoyl-CoA Hydratase in WNT. (a) Framework to find metabolic bottleneck genes using MBA and GPR. (b) Venn diagram showing unique and overlapped genes between Dawn WNT and Dawn 7h WNT. (c) Protein-protein interaction network of 28 unique bottleneck genes of Dawn 7h WNT indicates the regulatory role of nucleotide diphosphate kinase. (d) Venn diagram showing unique and overlapped genes between Dawn WNT and Dusk WNT. (e) Protein-protein interaction network of 38 unique bottleneck genes of Dusk WNT indicates the regulatory role of nucleotide diphosphate kinase and enoyl-CoA hydratase. (f) expression analysis of five different nucleotide diphosphate kinase genes.

Interestingly, gene ontology analysis revealed different functionality of bottleneck genes in different WNT conditions (Supplementary Fig. S4). In Dawn WNT condition, bottleneck genes are mostly from phosphorylation, small molecule metabolic process, and phosphate-containing compound metabolic process. Interestingly, in the Dawn 7h WNT condition, bottleneck genes are associated with cellular responses to oxidative processes, reactive oxygen species metabolic processes, and phosphorylation, among others. Conversely, for the Dusk WNT condition, bottleneck genes are linked to purine-containing compound metabolic processes, nucleobase-containing small molecule metabolic processes, and phosphorylation. Notably, phosphorylation emerges as a consistent top bottleneck process across all three WNT conditions, as revealed by gene ontology analysis. A previous study revealed that phosphorylation might play specific roles in amylopectin biosynthesis only in response to high-temperature stress for rice grain through different phosphorylation motifs such as sP, LxRxxs, and tP. As WNT conditions are metabolically distinct from each other, we next analyzed how metabolic bottlenecks dynamically progressed along the day compared to the Dawn WNT conditions.

Between Dawn WNT and Dawn 7h WNT conditions, there were 5 bottleneck genes that were unique to the Dawn WNT condition (Fig. 5 b). These are transcription initiation factor TFIID subunit 10, glutamine amidotransferase, cytidylyltransferase, pentatricopeptide, and urease. A gene ontology analysis revealed these genes are associated with nickel insertion and nickel cation binding. Nickel cation is a cofactor for certain enzymes, including urease, which is involved in nitrogen metabolism. A previous study (Liu *et al*., 2022) showed that nitrogen metabolism played spatial and temporal regulations in tissue (or stress)-specific way, indicating the importance of nitrogen metabolism in growth, development and stress responses. For the Dawn 7h WNT condition, there were 28 bottleneck genes that were unique to the Dawn 7h WNT condition (Fig. 5 b). A protein-protein interaction network revealed the central role of nucleotide diphosphate kinase in regulating bottleneck genes in Dawn 7h WNT (Fig. 5 c). Interestingly, nucleoside diphosphate kinase converts GTP to GDP, which is a key reaction in the ppGpp biosynthesis pathway. ppGpp is a stress regulator in many plants and bacteria (Hauryliuk *et al*., 2015). Between Dawn WNT and Dusk WNT conditions, there were 4 bottleneck genes that were unique to the Dawn WNT condition (Fig. 5 d). Apart from cytidylyltransferase, all other genes were similar to the unique Dawn WNT condition of Dawn WNT vs Dawn 7h WNT conditions, impacting nitrogen metabolism. For the Dusk WNT condition, there were 38 bottleneck genes that were unique to the Dusk WNT condition (Fig. 5d). A protein-protein interaction network revealed the central role of nucleotide diphosphate kinase (NDPK), like the Dawn WNT vs Dawn 7h WNT conditions (Fig. 5e). The rice genome contains five NDPKs. To determine the possible NDPK that may be responsive to WNT conditions in rice, we examined the expression of these five genes in an independent rice grain transcriptome dataset for control and WNT stress. Two of these genes, *OsNDPK1* and *OsNDPK4* were expressed strongly in rice grains relative to other gene family members (Fig. 5f). Expression of *OsNDPK4* is induced by the WNT stress, while transcript abundance of *NDPK1* decreased under WNT stress. Since NDPKs have a central role in energy utilization, a change in transcript abundance leading to differential enzymatic activity, could alter the ratio of carbon directed towards starch biosynthesis via ADP-glucose versus cell wall via UDP-glucose (Zeeman *et al*., 2010). Additionally, enoyl-CoA hydratase was also involved in regulating bottleneck genes in Dusk WNT. Enoyl-CoA hydratase is an important step in the β-oxidation, and interestingly it was shown for wheat that, up-regulation of genes belonging to fatty acid β-oxidation pathway (Aprile *et al*., 2013).

To understand the temporal WNT responsiveness of the bottleneck genes, we analyzed the expression of all 451 genes in the published transcriptome dataset generated from developing rice panicles treated with WNT stress along with the controls (Desai *et al*., 2021). Our clustering analysis revealed that expression of 67 genes was relatively high under WNT (Dawn 7h) compared to control conditions (Fig 6a). GO analysis of the cluster showed the enrichment of ATP related terms such as ‘ATP binding’, ‘ATP hydrolysis activity’, ‘ATPase complex’ and ‘ATPase-coupled transmembrane transporter activity’ (Supplementary Fig. S5). Among these genes, we found six subunits of ATP synthase and four ATP binding ABC transporters. Besides these genes, the cluster also consists of five vacuolar ATP synthases.

**Fig. 6.**
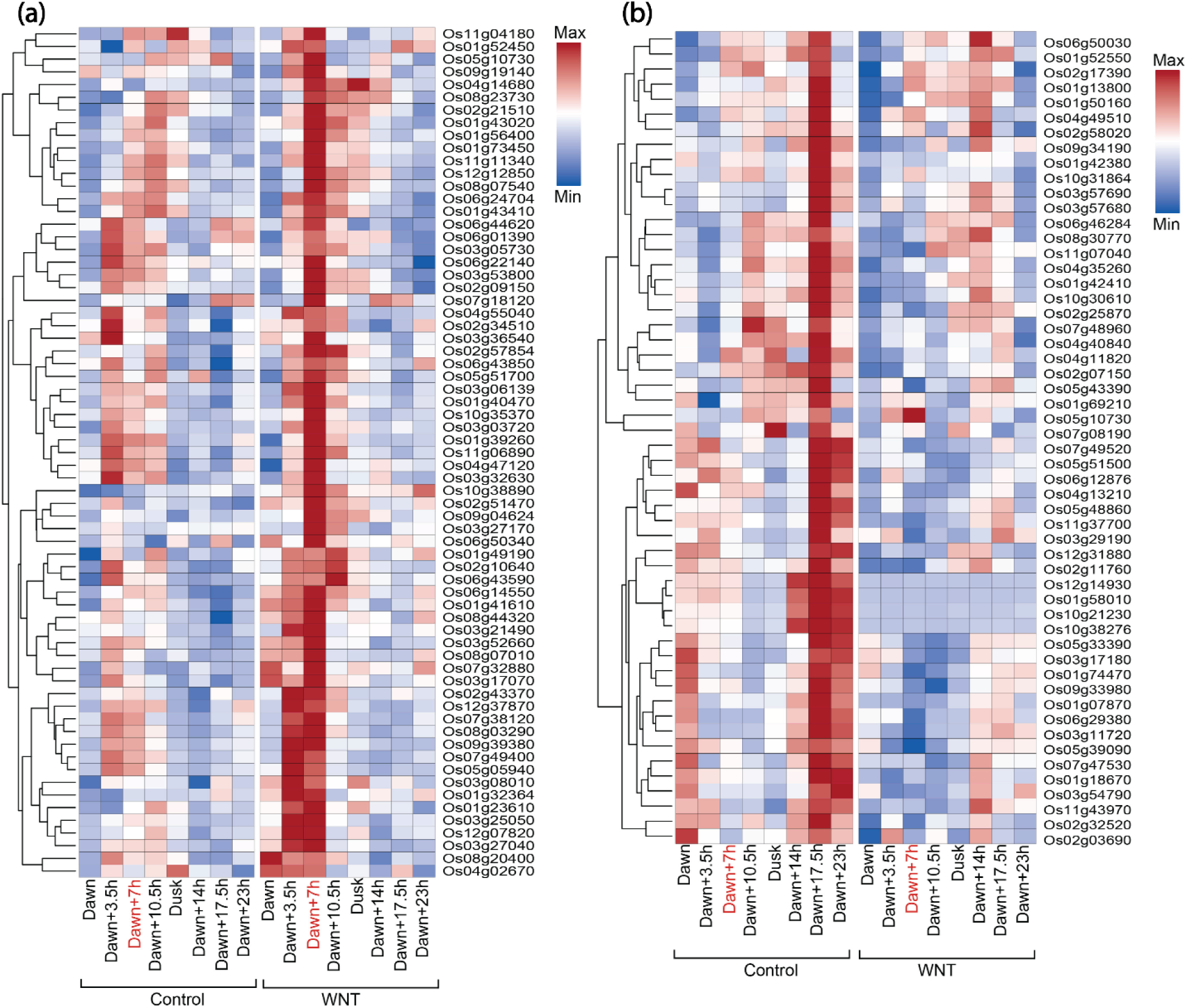
Two clusters derived from bottleneck genes showed highly similar responses to the temporal WNT treatment study. a) K-means clustering analysis showed that expression patterns of 67 genes are induced under WNT at Dawn+7 compared to the control conditions. b) Expression of 53 genes in suppressed in response to WNT at Dawn+17.5h.

ATP synthase complex, in particular vacuolar ATPase is known to play a role in response to oxidative stress, which indicates the WNT response role of genes in the cluster as the high temperature triggers cellular oxidative stress (Fortunato *et al*., 2023). For instance, the previous genetic study showed that an overaccumulation mutation in subunit E isoform 1 of the vacuolar H+-ATPase (*LOC_Os01g46980*) led to floury grain phenotype (Lou *et al*., 2021), wherein a similar developmental defect has also been observed in grain developed under WNT conditions (Dhatt *et al*., 2021). In addition to the genes induced by stress, 53 genes are found to be suppressed by WNT during the daytime (Dawn 17.5h) (Fig 6b). This cluster is primarily enriched in genes with a function such as ‘catalytic activity’, ‘ion binding’ and ‘small molecule binding’. Arabidopsis ortholog of two aldehyde oxidase coding genes (*LOC_Os03g57690* and *LOC_Os03g57680*) in the cluster catalyzes the abscisic acid (ABA) biosynthesis. Suppression of these ABA-related genes, a regulator of seed development and maturation, indicates that an imbalance in ABA concentration under WNT contributes to the distortion of seed development.

In summary, this study introduces the first-ever rice grain genome-scale metabolic model (GSM), iOSA3474-G, and investigates the impact of warmer night temperatures (WNT) on rice kernel quality, particularly chalkiness, due to global warming. Through synergistic computational-experimental approach, this work predicted markers like histidine and tyrosine linked to different growth phases under WNT conditions. Importantly, we have predicted MDAR and nucleoside diphosphate kinase as regulators of metabolic flux. These findings present new perspectives into understanding grain chalkiness in rice and lay the groundwork for innovative approaches to bolster crop resilience in the face of environmental challenges.

## MATERIALS AND METHODS

### iOSA3474-G Reconstruction

We used the maize kernel model from iZMA6517 (Chowdhury *et al*., 2023) as the template to reconstruct the iOSA3474-G. We downloaded the amino acid sequence for all the maize kernel genes from the MaizeGDB website. We collected amino acid sequence of all the genes for the entire IR-64 rice genome from the UGA Rice Genome Project (Buell, 2002). We then performed bi-directional blast between maize kernel genes from iZMA6517 and entire rice genome. Out of 6257 genes from the maize kernel, 3269 genes was matched with rice grain with an E-score 0.0, resembling high degree of homology between maize kernel and rice grain. The highest E-value for rice grain gene recorded was 9.5E-06 for *LOC_Os04g56710* and *LOC_Os02g11760* (Supplementary Data S3). Eventually, all the maize kernel genes from iZMA6517 has homologue in rice kernel. Moreover, the blast identified same rice grain genes for multiple maize kernel genes, thereby we automatically removed those duplicates rice grain genes from the model through an automatic python scripting. GPR for all the reactions were manually curated from the KEGG and Rice Genome Project database. As a result, 2899 out of 4060 reactions (71%) has GPR relationship. In addition, We collected expression data from literature (Desai *et al*., 2021) to see how many of those rice genes were expressed. If not expressed, those genes were discarded from the model. Next, we took a previously published rice grain model (Shaw and Cheung, 2021) and collected genes and reactions which were not part of the newly reconstructed draft model. For the genes, only those were added to the draft model which had expression values found in literature (Desai *et al*., 2021). For reactions, we considered those which were not part of the draft model. Subsequently, we determined compartments for those reactions using CELLO (Yu *et al*., 2006), and added to the draft model. GAMS, in conjunction with the CPLEX solver, was employed to tackle all optimization challenges. The NEOS server offers an alternative for executing GAMS codes without necessitating a license purchase. A comprehensive guide for running GAMS codes on the NEOS server is available in the literature (Chowdhury, Alsiyabi, *et al*., 2022). Additionally, to cater to the convenience of COBRApy users, an SBML version of the iOSA3474-G is provided.

### Expression Distributed Reaction Flux Measurement (EXTREAM)

The formulation of EXTREAM is the following:

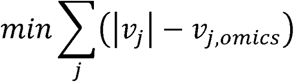

Subject to,

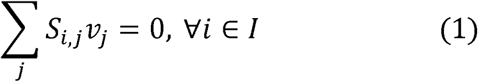

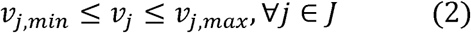

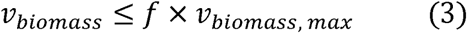

Here, *v_j_* is the flux to be calculated for reaction *J, v_j,omics_* is the reference condition calculated from the gene-protein-reaction association for reaction *J, s_i,j_* is the stoichiometric matrix for metabolite *I*, and reaction *J*, *v_j,min_* and *v_j,max_* are the upped and lower bound of reaction *J, v_biomass_* is the desired biomass growth rate, *v_biomass,max_* is the maximum possible biomass growth rate, and *f* is fraction between 0 to 1. To reformulation of EXTREAM algorithm to a linear optimization problem can be found in our previous work (Chowdhury *et al*., 2023).

### Parsimonious Flux Balance Analysis (pFBA)

pFBA (Lewis *et al*., 2010) is constrained based optimization technique to model GSMs. The pseudo-steady state mass balance in pFBA is represented by a stoichiometric matrix, where the columns represent metabolites, and the rows represent reactions. For each reaction, upper and lower bounds is imposed based on thermodynamic information. pFBA provides the flux value for each reaction in the model according by solving the following optimization problem:

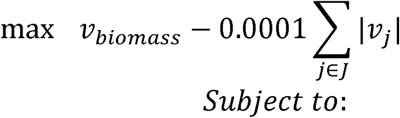

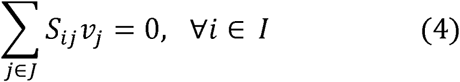

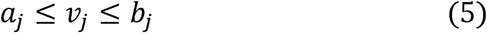

In this formulation, *I* is the set of metabolites and *J* is the set of reactions in the model. *S_ij_* is the stoichiometric matrix with *i* indicating metabolites and *j* indicating reactions, and *v_j_* is the flux value of each reaction. The objective function, *v_biomass_*, is the proxy of the growth rate of an individual cell. *a_j_* and *b_j_* are the lower and upper bounds of flux values for each reaction. For forward reactions, the highest possible bounds were 0 mmol/gDW/h to 1000 mmol/gDW/h. For the reversible reactions, the highest possible bounds were 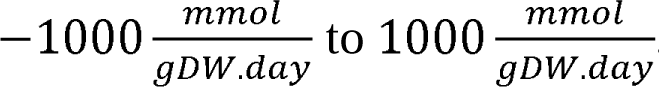.

### Metabolic Bottleneck Analysis

To determine the metabolic bottleneck in a GSM metabolic bottleneck analysis (Chowdhury *et al*., 2023) was used. The formulation is as following.

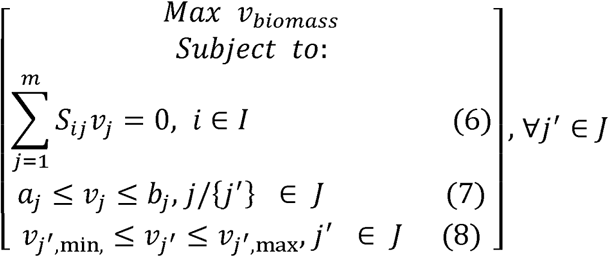

Here *a_j_* is the lower bound reaction *v_j_* and *b_j_* is the upper bound of reaction *v_j_*. Both *a_j_* and *h_j_* were calculated from the transcriptomics data and gene-protein-reaction association. *v_j_f_,min_* is the expanded lower bound of the reaction *j^’^* and *v_j_f_,max_* is the expanded upper bound of the reaction *j^’^*. In this case, we set 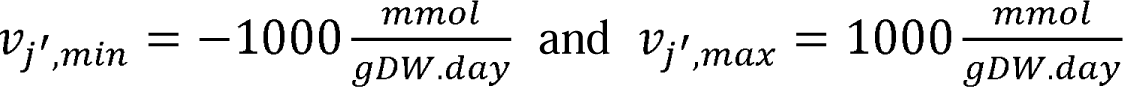. We solved the optimization problem by maximizing the biomass *v_biomass_* for the new expanded flux space of each reaction *j^’^* in an iterative manner and then recorded the biomass growth rate. From this biomass growth rate collections, we can check for which *j^’^* biomass growth rate increased significantly. Then that *j^’^* can be considered as the metabolic bottleneck of a given metabolic network.

### Grain Volume Measurement for Control and WNT Conditions

We used PI-PLAT imaging system to capture the RGB digital images of the primary panicles for the 3D reconstruction (Sandhu *et al*., 2019). In total, 120 images covering 360° angle were captured for each panicle from control and WNT treated plants using two digital cameras (Sony α6500) with LED as a light source (ESDDI PLV-380, 15 Watt, 5000 LM, 5600 K) moving on a wheel at a constant speed controlled by the eclectic motor system. In the image processing steps, digital RGB images captured by the PI-PLAT system were converted to hue saturation value space, and background objects such as the imaging chamber and wooden board were removed. These images were subjected to a denoising process to remove the remaining background. The pre-processed images were then used to construct a 3D point cloud of the panicle using the MVE pipeline. The generated point clouds were segmented, and scaling was performed to extract components of the panicle on a uniform scale by leveraging the positional information of the colors. Volume quantification was performed using voxelating the point clouds.

### Software and Hardware Resources

The General Algebraic Modeling System (GAMS) version 24.7.4 with IBM CPLEX solver was used to run pFBA, EXTREAM, and MBA algorithm on the model. Each of the algorithm was scripted in GAMS and then run on a Linux-based high-performance cluster computing system at the University of Nebraska-Lincoln. The model is also available in the systems biology markup language.

### SUPPORTING INFORMATION

Figure S1. WNT induced chalkiness in different rice genotypes.

Figure S2. Bottleneck genes in different conditions.

Figure S3. GO enrichment analysis for bottleneck genes in different control conditions.

Figure S4. GO enrichment analysis for bottleneck genes in different WNT conditions.

Figure S5. GO terms enriched among genes.

Data S1. Biomass composition of rice grain.

Data S2. Bottleneck genes in all conditions.

Data S3. Blast data between maize kernel and rice grain.

## ACKNOWLEDGEMENT

R.S. gratefully acknowledges funding support from the National Science Foundation (NSF) CAREER grant (1943310). H.W. acknowledges National Institute of Food and Agriculture (NIFA) Award #2024-67013-42385. The authors declare no competing interests.

## AUTHOR CONTRIBUTIONS

R.S. and H.W. designed the study and oversaw the project and funding acquisition; N.B.C. performed data curation, formal analysis, validation, and visualization; N.B.C. and A.N.C. worked on methodology; N.B.C. and A.N.C. wrote the original draft; R.S. and H.W. reviewed and edited the draft.

## DATA AVAILABILITY

The data that support for the findings of this study can be found in the related cited articles and/or in the supplementary data. All the codes used to generate these results can be accessed in the GitHub repository (https://github.com/ssbio/-rice_grain).

